# Generative adversarial networks as a tool to recover structural information from cryo-electron microscopy data

**DOI:** 10.1101/256792

**Authors:** Min Su, Hantian Zhang, Kevin Schawinski, Ce Zhang, Michael A. Cianfrocco

## Abstract

Cryo-electron microscopy (cryo-EM) is a powerful structural biology technique capable of determining atomic-resolution structures of biological macromolecules. Despite this ability, the low signal-to-noise ratio of cryo-EM data continues to remain a hurdle for assessing raw cryo-EM micrographs and subsequent image analysis. To help address this problem, we have performed proof-of-principle studies with generative adversarial networks, a form of artificial intelligence, to denoise individual particles. This approach effectively recovers global structural information for both synthetic and real cryo-EM data, facilitating per-particle assessment from noisy raw images. Our results suggest that generative adversarial networks may be able to provide an approach to denoise raw cryo-EM images to facilitate particle selection and raw particle interpretation for single particle and tomography cryo-EM data.

## INTRODUCTION

Despite the power of cryo-EM to determine new and challenging macromolecular structures, there remain a number of significant bottlenecks that continue to slow the adoption of cryo-EM and the throughput of structural data analysis. One such barrier is the inherent low signal-to-noise ratio (SNR) of individual single particle images due to the sensitivity of biological specimens to an incident electron beam (Conway et al., 1993; Glaeser, 1971; Henderson, 1995). The low SNR of the data is the result of the radiation sensitivity of biological samples, where even moderate electron doses of 40 - 60 e/Å^2^ still require dose compensation in order to remove the effects of radiation damage (Baker et al., 2010; Baker and Rubinstein, 2010; Grant and Grigorieff, 2015).

The low SNR of plunge-frozen macromolecules in cryo-EM datasets has led to the development of image analysis algorithms capable of dealing with uncertainty in single particle image alignment in order to determine 3D reconstructions. Currently, a number of software packages exist that are able to analyze hundreds of thousands of particles in order to reconstruct macromolecules at atomic-resolution such as RELION (Scheres, 2012), FREALIGN (Grigorieff, 2007), EMAN2 (Bell et al., 2016), cryoSPARC (Punjani et al., 2017), and IMAGIC (Afanasyev et al., 2017). These software packages typically apply a Wiener filter to a large number of particles from a range of defocus values in order to reconstruct an accurate representation of the 3D structure (Penczek, 2010).

Given the requirement for large dataset sizes, Singer and coworkers developed a per-particle image restoration approach - ‘Covariance Wiener Filtering’ - to restore information without the need of averaging images together (Bhamre et al., 2016). This work demonstrated that estimation of covariance matrix along with deconvolution could be used to perform amplitude correction of individual particles along in combination with denoising, thus highlighting a novel approach that could allow for per-particle assessment without 2D or 3D averaging of single particle images.

Looking beyond Covariance Wiener Filtering and Wiener Filtering, there are alternative approaches that could be used to denoise cryo-EM images, such as artificial intelligence. Within artificial intelligence, Generative Adversarial Networks (GAN) (Goodfellow et al., 2014) is a technique that combines two different neural networks together into a single pipeline - generative and discriminative neural networks - that could be used to recover information from images. The GAN involves training a discriminator to differentiate ‘good’ from ‘bad’ matches to a known training set, while also training the generator to alter the input images to better match the training set. GANs have been applied to a number of systems in order to recover information for more accurate interpretations. The ability of a GAN to recover information beyond the imaging resolution and noise limit was recently implemented successfully to recover faint and small-scale structures of galaxies (Schawinski et al., 2017). This implementation allowed the authors to recover low SNR images of galaxies after training GAN on a subset of high-resolution images of galaxies and then applied the resulting GAN on low SNR images of galaxies. Such an approach now offers astronomers the ability to recover features from slightly below the resolution limit of a given telescope using training from higher-resolution images of similar objects (e.g. galaxies).

Given the success of GAN on images of galaxies (Schawinski et al., 2017), we sought to explore its potential as a tool for denoising cryo-EM images. By using previously developed software for automated GAN parameter search and implementation on galaxy images (Li et al., 2017; Schawinski et al., 2017), we were able to recover structural information from synthetic cryo-EM data after training the GAN on single particle cryo-EM images. The GAN was capable of recovering information for structures that are closely resembling the training set, indicating that the GAN is capable of working on samples with similar structure. Finally, we demonstrated that GAN recovered information from experimental cryo-EM images, providing a proof-of-concept study for future applications of GAN to cryo-EM data analysis.

## RESULTS

In order to adapt GAN for cryo-EM, we utilized the previously described GAN software that was implemented for analyzing galaxy images (Schawinski et al., 2017). Given that this is already published, we will briefly summarize the GAN workflow here (Figure 1). The process for implementing GAN on cryo-EM data begins with training between model projections and particles. For synthetic data, models were derived from atomic cryo-EM structures or PDB models that were converted into cryo-EM density. The projections are then provided to the GAN alongside a ‘noise-added’ version of the projection. The GAN starts with the noise-added image and will begin generating different images. These images will be compared to the original projection via the discriminator, allowing the algorithm to optimize parameters that will restore noisy images to a high signal-to-noise ratio version, using the original projection as a guide.

**Figure 1:**
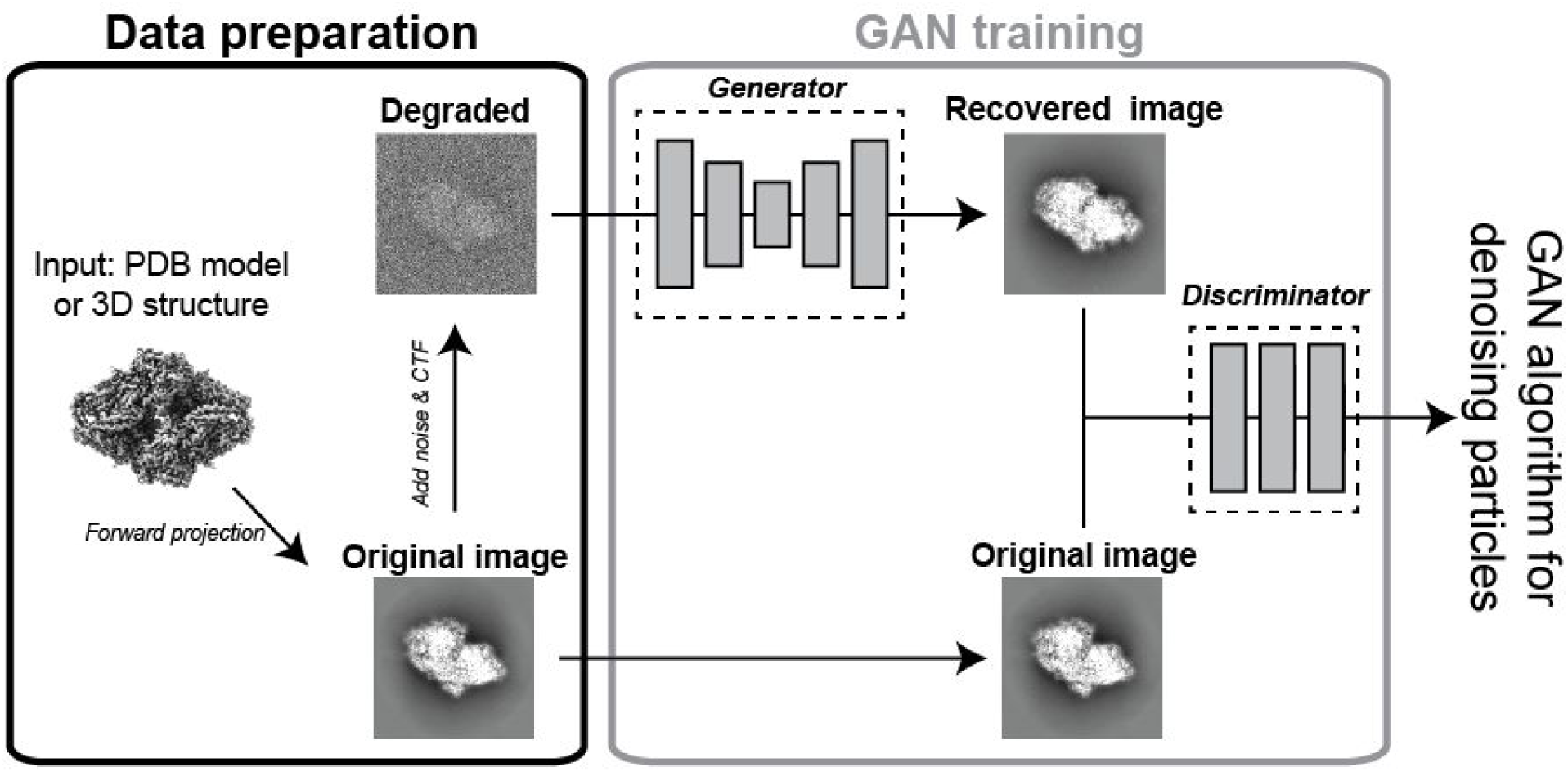
Schematic of GAN training for single particle image analysis. Forward projections of a 3D model were used as the ground truth to compare to noise-added synthetic data or real single particles. The ground truth projection then steers the generator algorithm to map the changes to the image required to convert the noise-added image into a higher SNR image. The discriminator is used to test the comparison between the recovered image and ground truth, with a final result being a ‘GAN-recovered’ image.

In order to test the feasibility of GAN to increase the SNR for cryo-EM data, we trained and tested projections of ß-galactosidase given its role as standard test specimen for cryo-EM. This test involved generating 2000 random projections of a 2.2 Å ß-galactosidase structure (Bartesaghi et al., 2015) and presenting them alongside images of the same projections, but with noise added. The resulting GAN was then tested using random projections from the same model with noise added, which resulted in a significant improvement in the relatie signal of the particle compared to the input noisy particles (Figure 2). Shown alongside the original projection, the GAN-recovered particle exhibits features that are consistent with the size and shape of ß-galactosidase, indicating that the GAN has recovered structural information regarding the underlying protein structure.

**Figure 2:**
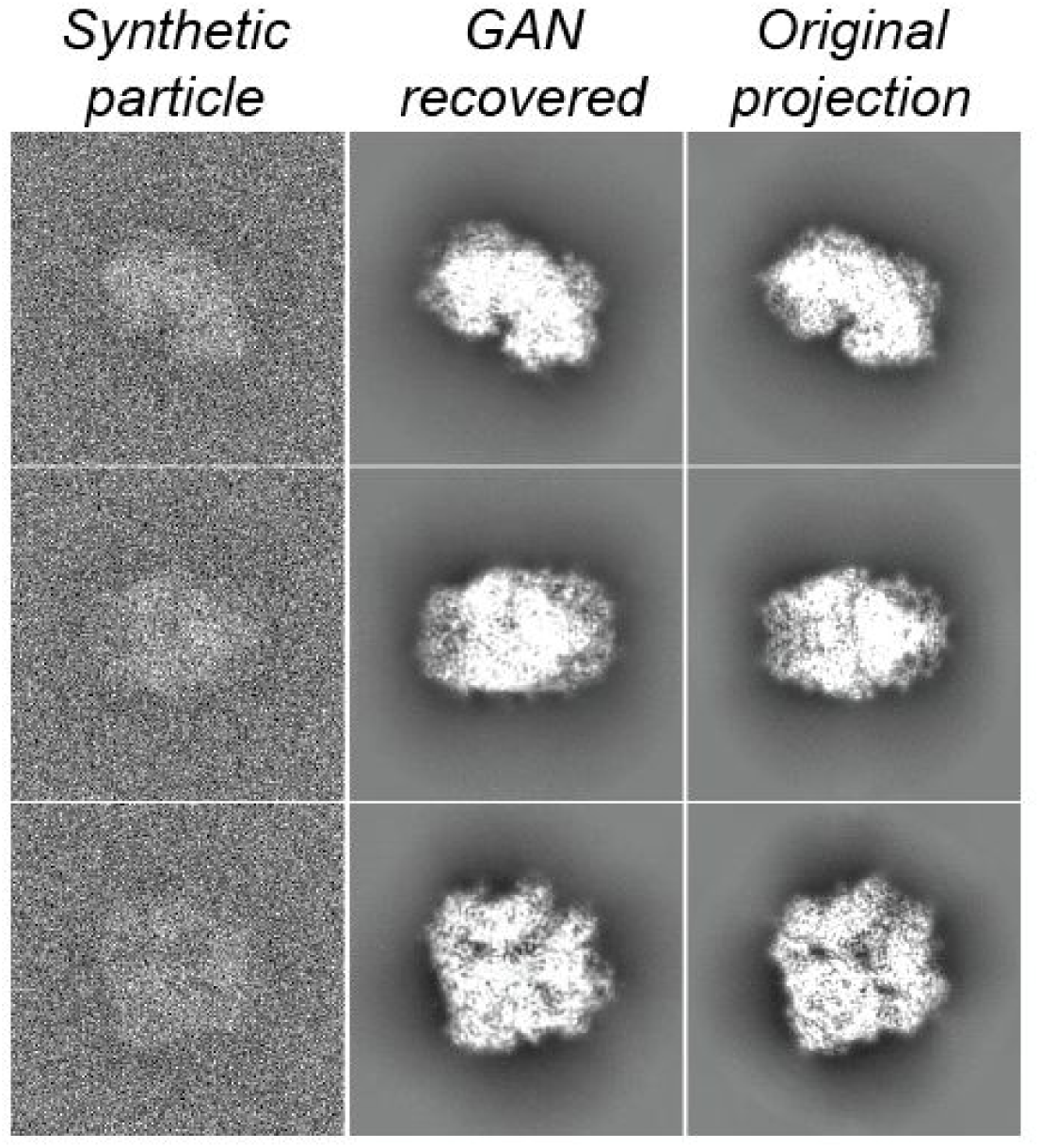
GAN recovers structural information from synthetic cryo-EM data. Shown are three different forward projections of ß-galactosidase (right, “Original projection”). For each projection, the ‘GAN recovered’ particle (center) was calculated by the GAN after training on 2000 pairs of original projections and noise-added projections (left).

To test the limits of GAN as a tool to recover information from single particle images, we created a panel of noise-added projections of ß-galactosidase. For each noise group, the GAN was trained in the same manner as the example shown above for Figure 2. After training, the GAN was then presented with random projections with noise-added particles containing the same level of noise as that used for the input projections (Figure 3). This demonstrated that the quality of GAN-recovered images was negatively impacted by increased levels of added-noise. For each of the views shown in Figure 3, the low SNR images highlight differences between the ground truth and GAN-recovered image. Despite the loss of structural features in the GAN-recovered image from the low SNR images, the overall shape of the particle remained mostly intact (Figure 3). This shows that the GAN is able to preserve low-resolution features of the particles, even when high-resolution features have been lost.

**Figure 3:**
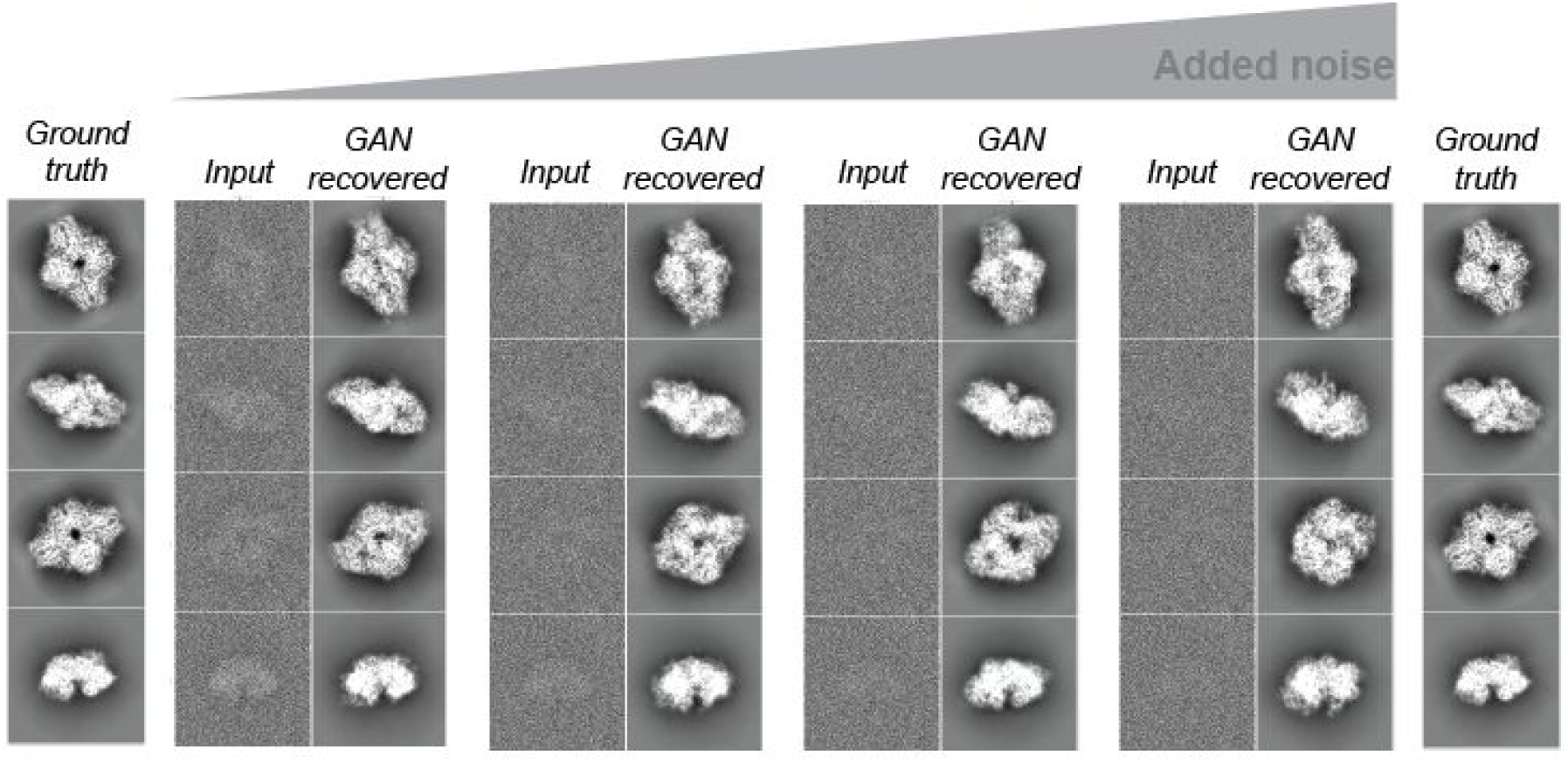
GAN recovers structural information from low SNR particle images. Increasing noise was added to projections of ground truth images (left and right), which was then used for GAN training. The resulting GAN was then tested on different images with the same noise, and the output image is shown ‘GAN recovered’.

These tests on ß-galactosidase suggested that GAN may represent a viable approach to recover structural features from cryo-EM images. To test this on a different biological system, we trained and tested a GAN on an atomic model of myosin V bound to an actin filament (Figure 4A) (Chen et al., 2012). While this atomic model was synthesized from extensive structural and biophysical studies, it remains an approximate model since there are no atomic structures of a single myosin V dimer on an actin filament. This is due to the challenging nature of the experiment, where the small size of myosin alongside sparse binding of along actin filaments results in a very difficult problem for particle picking.

**Figure 4:**
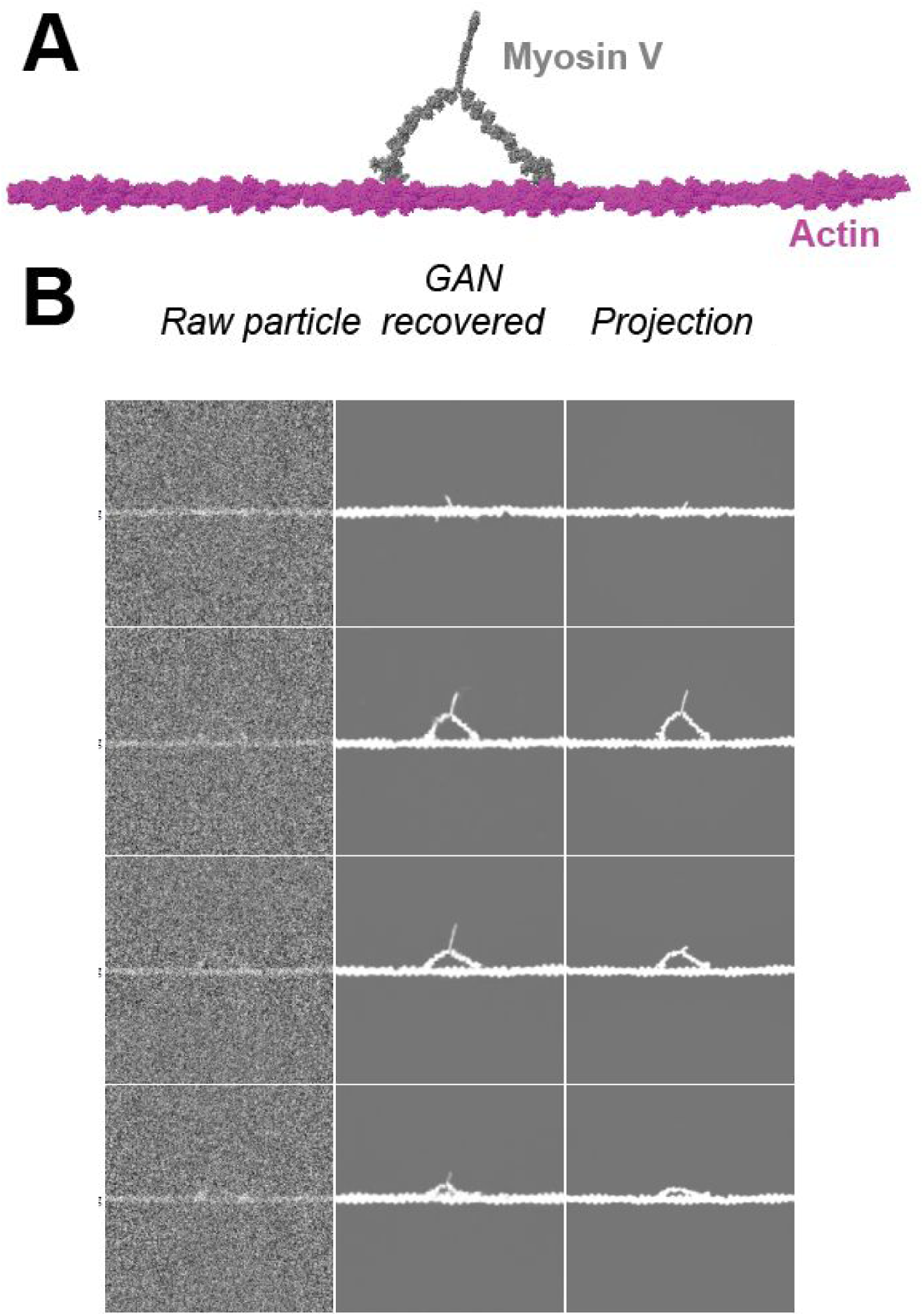
GAN denoises synthetic images of dimeric myosin V on actin filaments. (A) Atomic model of myosin V on actin filament. (B) Noise-added synthetic projections (left) are shown alongside GAN recovered images (middle) and the original projection of the model (right).

With this atomic model, we generated and tested the ability of GAN to recover the signal of myosin V motors on actin filaments (Figure 4B). The resulting GAN-recovered images showed that myosin V motors could be easily identified, whereas identifying the motor location in the raw, noisy particles involved more uncertainty. Importantly, as seen previously, the GAN-recovered image does introduce small features that differ from the original projection. This includes the angle of myosin motor domain relative to the actin filament, location of the tail, and angle of the tail relative to the filament. However, despite these differences, the GAN was able to successfully identify the individual motors on the actin filaments.

To further extend the potential applications of GAN for cryo-EM, we next turned to a different system - the binding of regulatory proteins to RNA Polymerase II (RNAPII). As the enzyme that templates the formation of mRNA from DNA, RNAPII is a large protein complex with many regulators that bind all across the surface in order to direct initiation, elongation, and termination of RNAPII activity (Nogales et al., 2017). While much has been discovered in recent years using cryo-EM there remain a number of key binding partners that have unknown binding sites on RNAPII and unknown mechanisms of action, requiring further structural studies. Thus, while the atomic structure of apo-RNAPII is known, there are many yet-uncharacterized structures of RNAPII bound to regulators.

With this framework in mind, we wanted to test whether a GAN trained on apo-RNAPII would be suitable for analyzing images of RNAPII bound to different regulatory binding partners (Figure 5A). To this end, we trained a GAN on projections of apo-RNAPII (PDB 1NT9) (Armache et al., 2003) using the approach described above. This GAN was tested on RNAPII-TFIIS complex (PDB 1Y1Y) (Cramer et al., 2004) and RNAPII-PIC (closed state) (PDB 5IY6) (He et al., 2016). These complexes were chosen to highlight the performance of the GAN on RNAPII-complexes for differing molecular weight as the size of TFIIS is < 5% of the total mass of RNAPII whereas the additional components that form the RNAPII-PIC (TBP, TFIIA, TFIIB, TFIIE, TFIIF, TFIIH, and DNA) are approximately the same size as RNAPII.

**Figure 5:**
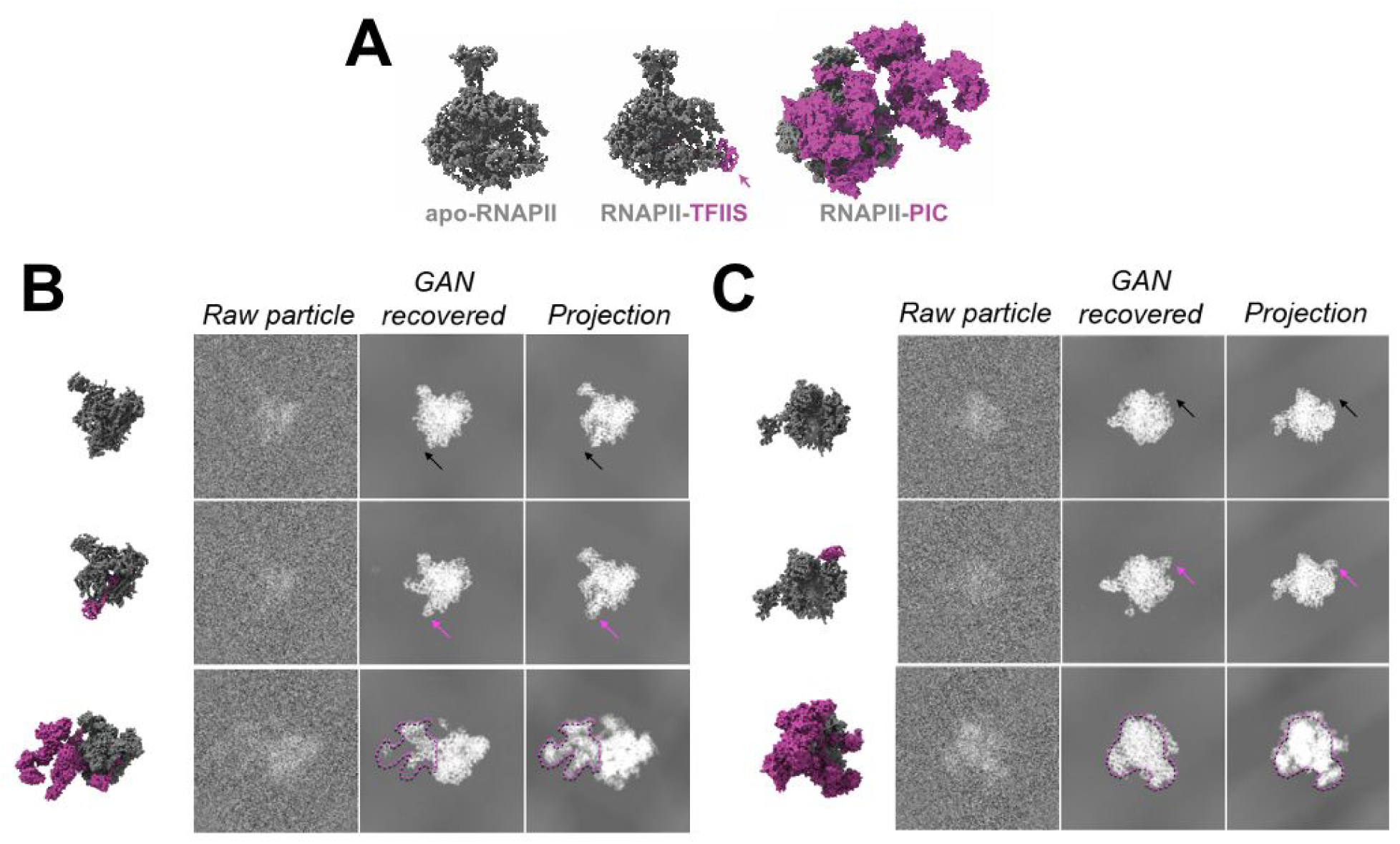
GAN training on apo-RNAPII allows recovery of information from higher-ordered RNAPII assemblies. (A) Atomic models of apo-RNAPII (PDB 1NT9), RNAPII-TFIIS (PDB 1Y1Y) and RNAPII-PIC (PDB 5IY6). RNAPII is colored in gray and additional binding partners (TFIIS, PIC) are colored in magenta. (B & C) (Top row) GAN trained on apo-RNAPII and tested on apo-RNAPII noise-added synthetic images. Black arrow indicates TFIIS binding location. (Middle row) GAN trained on apo-RNAPII and tested on RNAPII-TFIIS noise-added synthetic images. Magenta arrow highlights TFIIS density. (Bottom row) GAN trained on apo-RNAPII and tested on RNAPII-PIC noise-added synthetic images. Magenta dotted lines outline PIC density. (B) and (C) are two different viewing directions of RNAPII.

For all synthetic data projections, the GAN was capable of recovering overall structural details of the single particle (Figure 5). In the case of apo-RNAPII, the GAN denoised the projection so that the overall shape of RNAPII was recovered (Figure 5B&Figure 5C). For RNAPII-TFIIS, the GAN recovered images showed the presence of additional density that is located in the same location as where TFIIS interacts with RNAPII (Figure 5B & Figure 5C, pink arrow). However, as seen previously, there were differences observed between the recovered image and the ground truth, namely, the density of the stalk and jaw of the RNAPII appeared to be less prominent in apo-RNAPII whereas RNAPII-TFIIS had other densities that appeared in the GAN-recovered image (Figure 5B).

Finally, to test the limits of the GAN, we used the GAN trained on apo-RNAPII with particles that created from RNAPII-PIC, where the additional binding factors in the PIC comprise approximately the same size of apo-RNAPII. The GAN recovered images from RNAPII-PIC showed the presence of a additional density (Figure 5B & Figure 5C, pink outline) that comprised some (Figure 5B) or all (Figure 5C) of the additional density expected for the PIC. Therefore, there comparisons highlight the ability of GAN to recover structural information from simulated cryo-EM images of structures that had not be previously trained with the GAN.

Next, we wanted to test the performance of GAN with experimental cryo-EM data. This was done by training and testing the GAN with the 2.2 Å ß-galactosidase dataset (Bartesaghi et al., 2015), where polished, refined particles from the final iteration of RELION auto-refine were included with forward projections of the final reconstructed map (without sharpening) for GAN training (Figure 6A). Following this training, the resulting GAN was tested on other particles that were previously not analyzed by the GAN (Figure 6B). This test demonstrated the ability of the GAN to recover structural information from the raw cryo-EM particles, as comparison of the GAN recovered image to the original projection highlights the overall similarity of the images. Interestingly, we observed a small number (<1%) of GAN output images to show a different image than the projection image from RELION (Figure 6C). While we do not know the ground truth for this example, we speculate that the GAN might have assigned the correct euler angle, although further work will be needed to address this.

**Figure 6:**
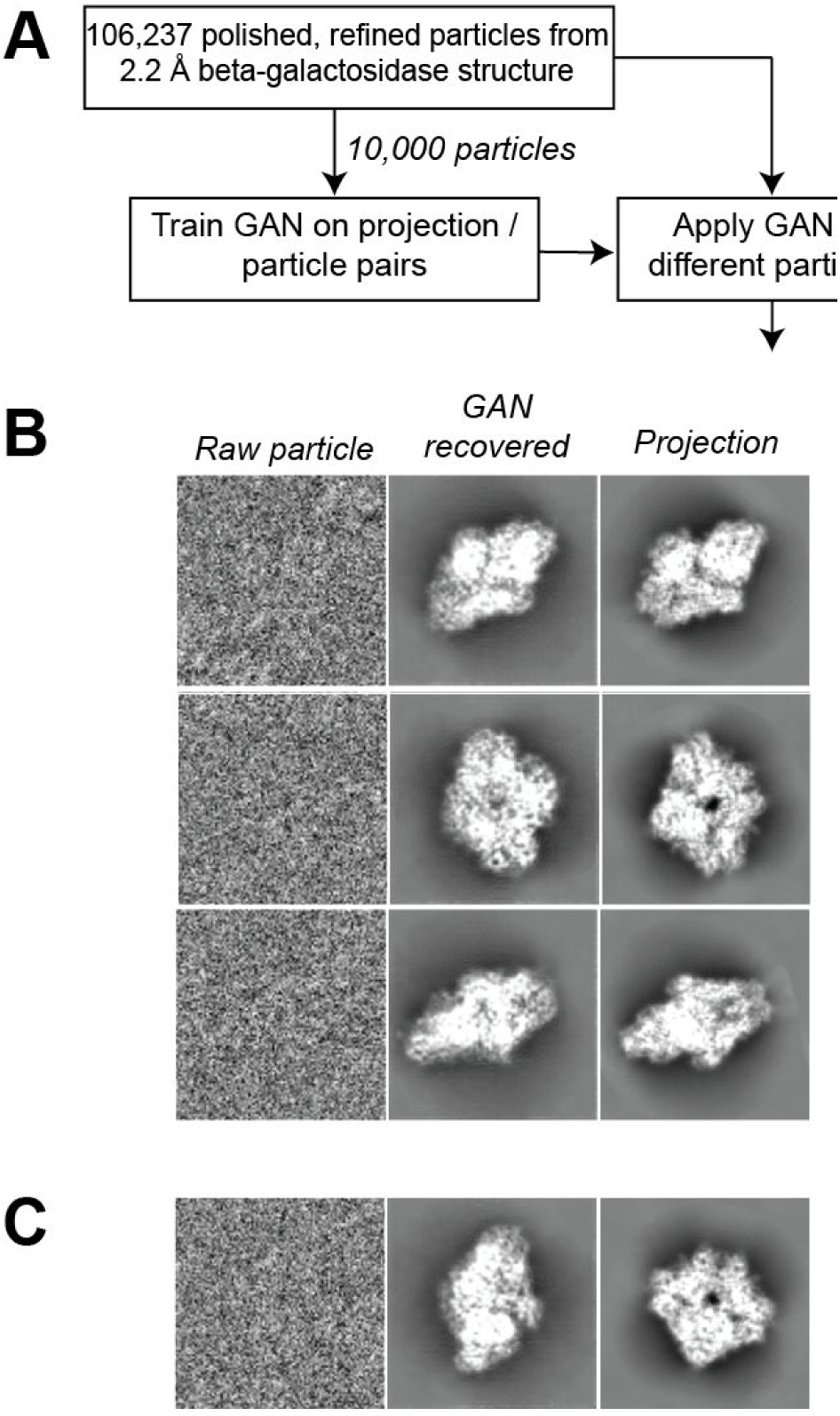
Training and implementation of GAN on experimental cryo-EM data of ß-galactosidase. (A) Outline of GAN training and testing for real ß-galactosidase particles. Using 10,000 polished, aligned particles from a 2.2 Å RELION refined structure of ß-galactosidase, forward projections of the 2.2 Å density was used alongside the raw particles for GAN training. This GAN was tested using different particles not previously analyzed by the GAN and the results are shown in (B). (B) Raw particles (left) are shown alongside the output GAN recovered images (middle) and the original projection (right). (C) <1% of GAN-recovered images did not match the original projection.

## DISCUSSION

This work provides a proof-of-concept study that highlights the ability of GAN, a form of artificial intelligence, to recover structural information from both synthetic and real cryo-EM single particles. In all cases, the GAN was trained on specific structures as training sets that comprised identical or similar samples, indicating that prior knowledge was required before the GAN could be implemented. From these comparisons, we observed that GAN is capable of recovering information while retaining the overall structural features. Importantly, the GAN was able to recover information from real cryo-EM data.

Despite the ability of GAN to recover structural information from cryo-EM data, the GAN did introduce differences in the recovered images that were not present in the original projections. This observed difference between the GAN output and the ground truth likely results from the low SNR of cryo-EM data; the GAN is capable of recovering the overall shape and structure, internal details are imprecise. Further work will be needed to overcome this significant limitation of the way that we implemented the GAN on cryo-EM data.

The results presented here indicate that GAN may be able to provide useful information for assessing individual particles during particle picking or protein complex integrity. Since the low resolution information is retained in the GAN recovered image, it is likely that the GAN will be able to identify particles in a cryo-EM micrograph. Moreover, beyond particle localization, the GAN may be able to return information to the user regarding the state of the protein complex if the structure of the protein is previously known. This suggests that GAN may help to increase the throughput of cryo-EM if raw micrographs can be used for sample assessment.

This work represents an important first step in the implementation of artificial intelligence-driven denoising of cryo-EM data. However, while we believe the GAN is recovering global structural information, we realize that much more work is necessary to understand how to implement GAN effectively for cryo-EM. In particular, we believe that alternative training strategies might be able to steer GAN parameter search towards high-resolution image recovery. These changes, in conjunction with abilities to test and prevent overfitting (i.e. “Einstein from noise” (Henderson, 2013)), may help to develop GAN into a part of the cryo-EM workflow.

## ACKNOWLEDGEMENTS

We would like to thank Zev Bryant (Stanford University) for the atomic model of myosin V-actin. K.S. acknowledges support from Swiss National Science Foundation Grants PP00P2_138979 and PP00P2_166159 and the ETH Zurich Department of Physics. CZ and the DS3Lab gratefully acknowledge the support from the Swiss National Science Foundation NRP 75 407540_167266, IBM Zurich, Mercedes-Benz Research&Development North America, Oracle Labs, Swisscom, Zurich Insurance, Chinese Scholarship Council, and the Department of Computer Science at ETH Zurich, the GPU donation from NVIDIA Corporation, the cloud computation resources from Microsoft Azure for Research award program.

## METHODS

### GAN training and testing on synthetic and real cryo-EM projections

For synthetic data, noise-added training data was generated by forward projections of 3D models using RELION (Scheres, 2012) with noise added using sigma=200. MRC-formated projection images were then processed on Titan-X GPUs using standard pix2pix pipeline that translates one image into another (https://github.com/phillipi/pix2pix). For real cryo-EM data, the output particles from RELION polishing and 3D auto-refine of the 2.2 Å ß-galactosidase structure (Bartesaghi et al., 2015) were presented to the GAN alongside forward projections of the density map without sharpening generated by RELION.

